# Assembled Cell-Decorated Collagen (AC-DC) bioprinted implants mimic musculoskeletal tissue properties and promote functional recovery

**DOI:** 10.1101/2021.06.22.449431

**Authors:** Kyle W. Christensen, Jonathan Turner, Kelly Coughenour, Yas Maghdouri-White, Anna A. Bulysheva, Olivia Sergeant, Michael Rariden, Alessia Randazzo, Andrew J. Sheean, George J. Christ, Michael P. Francis

**Author notes:** These authors contributed equally to this work (Kyle W. Christensen, Jonathan Turner). Senior authors (George J. Christ, Michael P. Francis).

## Abstract

Musculoskeletal tissue injuries, including the damage and rupture of ligaments and tendons, and volumetric muscle loss (VML), are exceptionally commonplace and often lead to permanent disability and deformation. We developed an advanced biomanufacturing platform producing cellularized collagen microfiber implants to facilitate functional repair and regeneration of musculoskeletal soft tissues. This Assembled Cell-Decorated Collagen (AC-DC) bioprinting process rapidly and reproducibly forms 3D implants using clinically relevant cells and strong, microfluidic extruded collagen fibers. Quantitative analysis showed that the directionality and distribution of cells throughout AC-DC implants mimic the cellular properties of native musculoskeletal tissue. AC-DC bioprinted implants further approximate or exceed the strength and stiffness of human tendons and ligaments and exceeded the properties of commonplace collagen hydrogels by orders of magnitude. The regenerative potential of AC-DC implants was also assessed *in vivo* in a rodent VML model. A critically sized muscle injury in the hindlimb was created and repaired, and limb torque generation potential was measured over 12 weeks. Both acellular and cellular implants were found to promote functional recovery compared to the unrepaired group, with AC-DC implants containing therapeutic muscle progenitor cells promoting the highest degree of recovery. Histological analysis and automated image processing of explanted muscle cross-sections revealed increased total muscle fiber count, median muscle fiber size, and increased cellularization for injuries repaired with cellularized implants. These studies introduce the tremendous potential of an advanced bioprinting method for generating tissue analogs with near-native biological and biomechanical properties with the potential to repair numerous challenging musculoskeletal injuries.

**One Sentence Summary:** Bioprinted collagen microfiber-based implants mimic musculoskeletal tissue properties *in vitro* and promote functional recovery *in vivo*.

## INTRODUCTION

Musculoskeletal tissue injuries are exceptionally commonplace in athletes, military, civilians, men, and women, from young to old, with over 30 million injuries reported annually in the United States alone *(1)*. For warfighters, extremity injuries account for up to 70% of trauma cases in theater *(2)*. These injuries, including the damage and rupture of ligaments and tendons, tears at the myotendinous junction (MTJ), and volumetric muscle loss (VML) have been observed among military service members and civilians alike as sequelae of high energy trauma and recreational activity. These traumatic injuries often require surgical intervention as they exceed the body’s self-repair capabilities and can significantly delay or entirely prevent a return to regular activity. Tendon and ligament injuries often require surgery, with as many as 250,00 repairs of the rotator cuff *(3)* and 200,000 reconstructions of the anterior cruciate ligament (ACL) *(4)* performed annually. As the most abundant tissue in the human body, comprising ∼40% of total body mass *(5)*, skeletal muscle damage also frequently occurs, with as many as 55% of sport injuries involving damage at the myofiber level *(6)*. While muscle has a high regenerative capacity for relatively minor injuries, natural repair mechanisms are overwhelmed in cases of VML and lead to chronic functional deficits *(7–10)*.

Unfortunately, the effectiveness and long-term outcomes of current traumatic musculoskeletal soft tissue treatment options, particularly with severe muscle involvement, are often limited *(11)*. Rotator cuff repair, for example, has been reported with re-injury rates greater than 50% *(12)*, can have coincident muscle damage, and often leads to permanent stiffness and limited range of motion. Tendon- and ligament-focused therapies, such as autografting, can lead to loss of function and morbidity at the donor site *(13)*, and cadaveric grafts can be subject to complications due to tissue availability and immunoreactivity. Treatment options for VML are currently severely limited and include autologous free muscle flap transfer *(7, 14)*, muscle transposition *(7, 8)*, or amputation and power bracing *(14)*. Again, donor site morbidity, when tissue is available for harvest, and the need for a uniquely skilled surgical team decrease positive patient outcomes and complicate VML treatment *(14)*.

Numerous additive biomanufacturing, or three-dimensional (3D) bioprinting, approaches have been explored to produce biological scaffolds and tissue mimetics with regenerative potential *(15, 16)*. Toward addressing VML in particular, several tissue engineering approaches have been investigated to restore function to injured muscle *(17–24)*. Additive biomanufacturing offers the potential for on-demand creation of clinically relevant implants capable of facilitating the restoration of functional native-like tissue. Thus, it has the potential to improve current treatment paradigms for musculoskeletal injuries significantly *(25)*. A biomimicry approach is often favored, where recreating the biochemical, morphological, and functional properties of targeted tissue is paramount to the therapeutic utility of manufactured implants *(15)*. In addition to mimicking the structural and material properties of native tissue, therapeutic cells are often included as key components of implants. Cell-based treatments offer the potential to improve the treatment of degenerative, inflammatory, and traumatic musculoskeletal disorders *(26)*, and cellularized constructs may promote more rapid and complete tissue regeneration compared to the use of biomaterial scaffolds alone.

However, to date, biomanufacturing approaches incorporating cells are often unable to adequately recreate fundamental properties of native tissue, limiting their use in a clinical setting. For example, soft hydrogels are the primary structural component used in typical bioprinting approaches *(27)*. Yet, they fail to match the mechanical properties of native tendon and ligament tissues by orders of magnitude *(28–30)*. In efforts to improve the strength of hydrogel-based or even cell or ECM-only constructs, bioreactors and extended lengths of *in vitro* culture have been used to facilitate cellular remodeling and self-assembly *(31–33)*. While this has led to marked improvements in strength and histological likeness to native tissue, the approach is costly, may have limited scalability commercially, and the produced scaffolds ultimately still fail to match the mechanical properties of native tissues. Specialized hybrid bioprinting approaches have been developed that incorporate thermoplastic polymers and hydrogels to improve the mechanical properties of printed parts with a modest increase in biological activity *(34, 35)*. However, composite scaffolds’ mechanical and biological properties may still differ significantly. Further, synthetic materials often elicit a chronic inflammatory response and limit tissue remodeling and regeneration *in vivo*.

As an alternative to traditional 3D bioprinting, various fiber-based biomanufacturing approaches have been explored *(36)*. These approaches build on the well-established clinical use of textiles and textile manufacturing processes, with additional means to seed scaffolds with cells after fabrication. Of particular importance for musculoskeletal tissue repair, synthetic *(37, 38)* and natural biomaterial fibers *(39–42)* may offer mechanical properties approximating native tissue *(40, 41)*. Fibrous biomaterial scaffolds have been directly produced by wet-spinning *(42, 43)*, electrospinning *(44, 45)*, and direct writing *(46)*, in which fiber production by solvent evaporation, polymerization within a solution bath, or temperature-based recrystallization, respectively, are integral aspects of the scaffold fabrication process. Alternatively, fibers can be produced as a feedstock for secondary assembly processes *(36)* by wet spinning, microfluidic spinning, biospinning, interface complexation, or melt spinning before being assembled into designed geometries as biotextiles by weaving *(37)*, knitting *(38)*, braiding *(41)*, or manual assembly *(40)*. In addition, across varying fabrication approaches, scaffolds have been seeded with cells after fabrication by manual pipetting, casting, and perfusion-based cell seeding, among other techniques *(36)*.

The assembly of biomaterial fibers into cellularized 3D geometries has relied almost exclusively on complex manual assembly by weaving, knitting, braiding, or winding *(36)*. These approaches offer severely limited reproducibility and scalability, both critical concerns for transitioning biomanufacturing processes and produced implants toward commercial scale and clinical use *(16)*. More automated and well-characterized fiber-based approaches, such as electrospinning, have seen success clinically. However, these scaffolds typically still fail to match native tissue biological or mechanical properties and commonly require cytotoxic processing conditions during manufacturing, precluding the controlled use of therapeutic cells in-process. Post-fabrication manual cell seeding has been used to cellularize such implants, but seeding processes typically lack precision, control, feedback, are subject to human variability, and may depend on the micro- and macro-scale scaffold geometry. Controlled cell seeding remains a significant challenge for scaffolds with complex geometries, high thicknesses, low porosity, and especially for producing scaffolds with designed and controllable cellularity throughout. Approaches that coat natural and synthetic threads with cell-laden hydrogels have been developed to avoid the need for manual cell seeding but again require a complex manual assembly, yet with limited control and reproducibility to form 3D structures *(47, 48)*. Due to the inherent variability of manual assembly and cell-seeding processes, the development of automated, controlled, and consistent biomanufacturing processes is critical for producing clinically relevant cellularized implants.

In this study, advanced bioprinting strategies that control for scalable, high output, and high print-fidelity processes are developed, where clinically relevant cells are positioned controllably along clinically relevant, high strength collagen fibers to biomanufacture musculoskeletal tissue analogs for restoring form and function to injured tissues. Human mesenchymal stem cells (hMSCs) or rat muscle progenitor cells (MPCs) are printed, targeting tendon- and ligament-focused or VML-focused applications, respectively. Mesenchymal stem cells offer excellent potential for augmenting musculoskeletal tissue repair and regeneration due to their immune-evasive properties *(49, 50)*, therapeutic effects *(50–52)*, multilineage differentiation potential *(53)*, and availability as a commercial clinically relevant cell type. Similarly, MPCs have shown marked therapeutic effects in facilitating functional recovery in volumetric muscle loss injuries in validated animal models *(11, 54)*. This study is the first report of using actual collagen fibers with high tensile strength as a filament for bioprinting and is a further significant advancement across the 3D biofabrication field by recreating the structural, cellular, and mechanical likeness of native tissue in an automated, scalable fabrication process, which was previously an ambitious and unrealized challenge *(15, 16)*.

## RESULTS

### Subhead 1: Bioprinting concept and design

We established an original Assembled Cell-Decorated Collagen (AC-DC) bioprinting method (Fig. 1) to produce implants approximating native musculoskeletal tissue properties. AC-DC bioprinting controllably seeds or “decorates” cells onto collagen microfiber as it passes through a cell-seeding reservoir before being wrapped next to and on top of itself to produce aligned 3D constructs mimicking native, regularly organized (aligned) musculoskeletal tissue architecture (movie S1). Printed implants are formed on temporary rigid frames, which maintain fiber alignment and 3D macrostructure during printing. With AC-DC bioprinting, fiber-based implants with designed width, length, thickness, and porosity, based on the print bed conformation and the computer-aided design (CAD) file, can be rapidly and repeatably produced via a fully automated process.

**Fig. 1.**
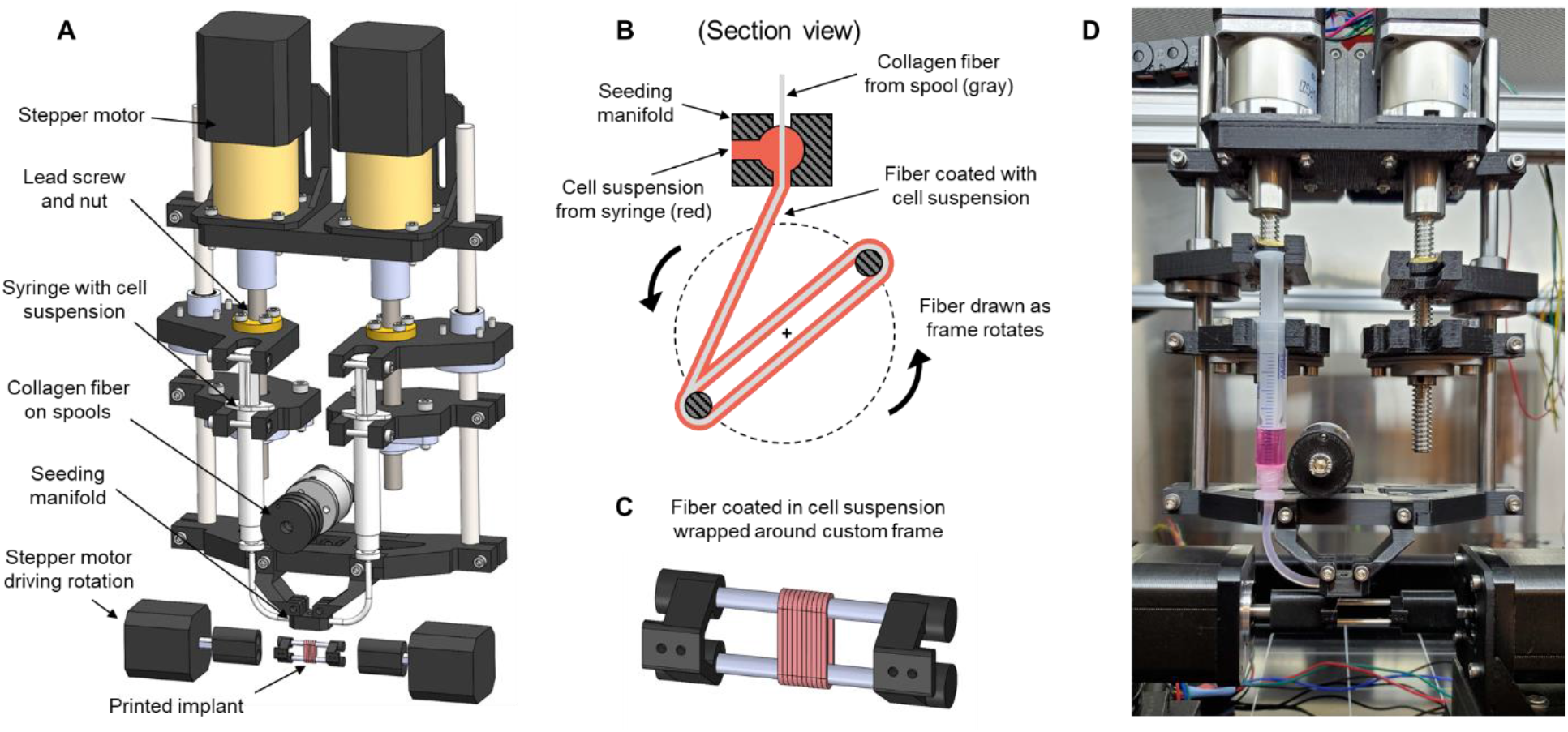
Assembled Cell-decorated Collagen (AC-DC) bioprinting concept and design. (**A**) Computer-aided design (CAD) model of our custom extrusion printhead and collection assembly with critical components identified. (**B**) Section view illustrating the cell seeding and implant biofabrication process. Collagen fiber passes through a seeding manifold, where it is uniformly coated with a cell suspension. The coated fiber is drawn onto a frame of arbitrary shape and dimensions and wrapped in parallel and on top of itself to form a 3D implant. (**C**) CAD model of a parallel fiber implant formed on a rigid frame with a multifunctional design. Frames maintain implant fidelity during culture, and implants are secured with sutures and easily removed from frames before further testing or implantation. (**D**) Photograph of printhead and collection assembly housed in a laminar flow hood during a single cell-solution bioprint.

A custom extrusion printhead (Fig. 1A) was designed and incorporated with a Folger Tech FT-5 R2 commercial 3D printer. Two separate planetary geared stepper motors with lead screw assemblies mechanically compress disposable syringes individually, extruding cell suspension with sub-microliter resolution. Multiple cell-solution printheads can prospectively facilitate the production of heterogeneous implants with differing cell populations in distinct regions (Fig. S1). Cell suspensions prepared in hyaluronic acid (HA) solutions, acting as “cellular glue”, were extruded into the seeding manifold during printing, where cells uniformly coated collagen fiber being drawn through the manifold (Fig. 1B). The volume of cell suspension extruded per millimeter of drawn fiber is a user-determined process parameter, offering a means to control the resulting cell density and the total number of cells throughout an implant. A custom printed implant collection assembly was designed (Fig. 1A) in which small rigid frames (Fig. 1C) are held between two stepper motors. Typically, frame designs consist of two horizontal bars or dowels kept a fixed distance apart by custom 3D-printed frame ends, forming a central opening to limit cell migration from the implant to the frame and enable improved nutrient diffusion to the implant from the surrounding cell culture media.

### Subhead 2: Implant fidelity and cellularity

Cellularized collagen microfiber implants were printed rapidly and repeatably onto various frame geometries (Fig. 2A-C) with centimeter-scale implants produced in minutes and no perceived loss of fidelity after hours of consecutive printing. After securing implants into bundles using sutures, they were easily removed from a frame (Fig. 2D) by removing one or both PLA frame ends and sliding implants off the steel dowels. Across varying custom geometries, rigid frames were found to maintain fiber alignment and the macrostructure of printed implants in culture over several weeks. Transmitted light microscopy of printed implants shows striations indicative of densely packed parallel fiber after four days of culture (Fig. 2E). Fluorescence imaging of implants printed with hMSCs stained with a cytoplasmic label (green), dead cell nuclear label (red), and fiber autofluorescence (blue) after four days of culture shows cells distributed throughout (Fig. 2F). Notably, cells were visibly elongated parallel to the collagen fiber direction, as is characteristic of native tendon and ligament histology. A grouping of three collagen microfiber strands (printed simultaneously from 3 spools of fiber attached to the printhead) was printed with hMSCs and labeled to assess the precision of the printing process (Fig. 2G). These three fiber strands act as the “building blocks” of printed implants, as three fiber strands are continuously drawn through the printhead, seeded with cells, and wrapped next to and on top of one another to form 3D implants.

**Fig. 2.**
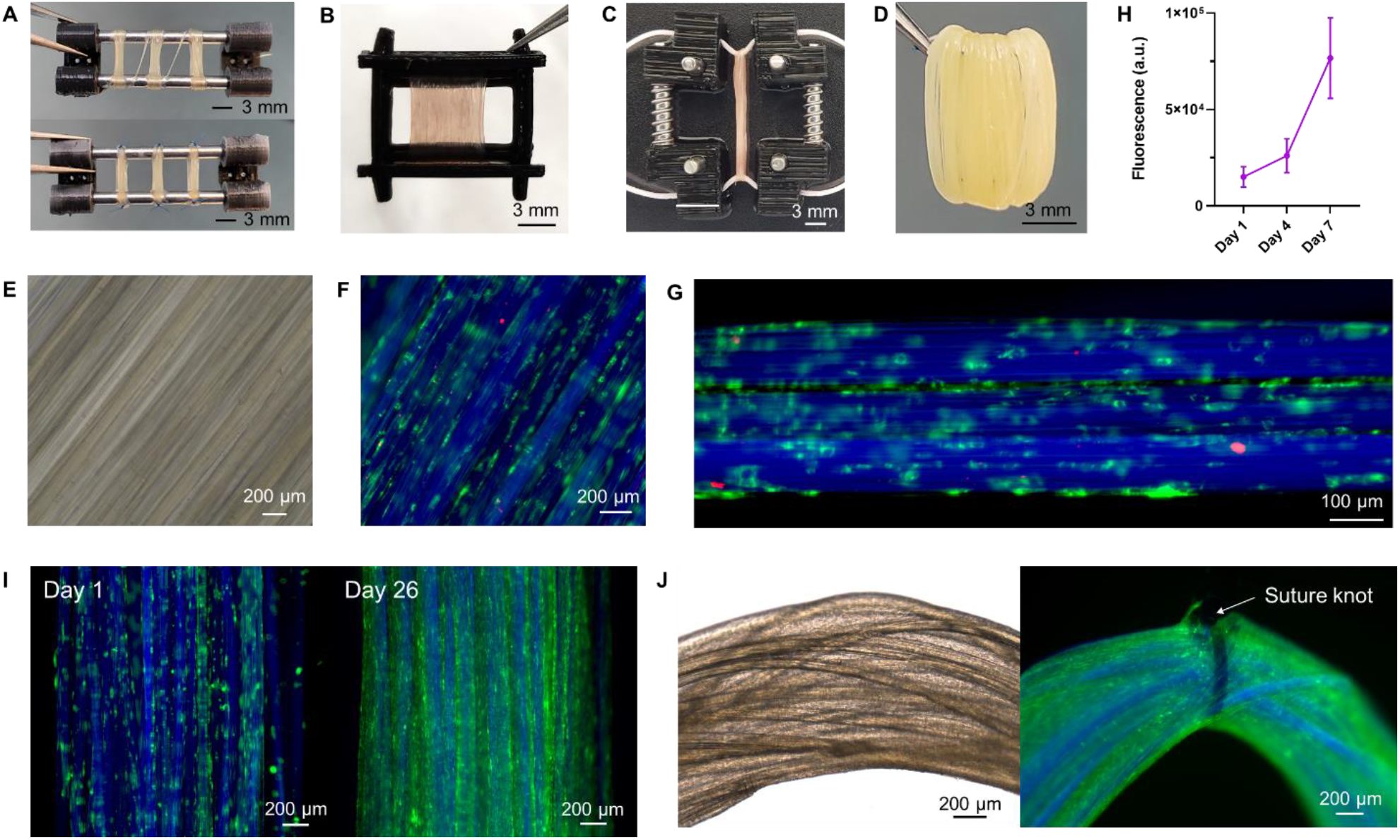
Implant fidelity and cellularity. (**A**) Three printed implants on a multifunctional frame before (top) and after (bottom) being secured with suture. (**B**) A larger printed implant on a custom frame geometry. (**C**) An implant is printed onto two lengths of sutures and held in tension by a custom frame. (**D**) Tweezers hold a printed implant after being secured with a suture and removed from a frame. (**E**) The transmitted light image of a printed implant showing striations indicative of densely packed parallel fiber after four days of culture. (**F**) Fluorescence image showing hMSCs distributed throughout and elongating parallel to the fiber direction after four days of culture. Fluorescence images F-I show all cells with the cytoplasmic label DiD (green) and collagen fiber autofluorescence at 405 nm (blue), with images F and G additionally showing dead cell nuclei with EthD-1 (red). (**G**) Fluorescence image showing hMSCs attached to and distributed along three “building block” strands of collagen fiber after four days in culture. (**H**) Fluorescence intensity indicating cell metabolic activity using the alamarBlue assay for implants printed with hMSCs after 1, 4, and 7 days of culture (n=6). (**I**) Fluorescence images of a printed implant showing an initial distribution of cells after 1 day of culture and a confluent densely-cellularized implant after 26 days of culture. (**J**) Transmitted light image (left) and fluorescence image (right) of implants secured by suture showing dense cellular ingrowth of hMSCs between and on top of collagen fibers after 6 weeks of culture.

Cell viability throughout printed implants was assessed qualitatively and quantitatively by fluorescent imaging of hMSCs. Qualitatively, generally, high viability is indicated by live cells (green), greatly outnumbering dead cells (red) (Fig. 2F, G). Quantitatively, ImageJ was used with established cell counting techniques to compare the number of live and dead cells throughout implants immediately after printing. For representative implants printed with typical process parameters, hMSCs were found to be 93.2±1.7% viable immediately after printing, and cell viability was consistently above 90% for various implants geometries and printing conditions. However, quantifying viability by fluorescent imaging becomes difficult beyond several days of culture, as cells become confluent throughout the implants, with individual cells becoming indistinguishable. As such, benchmark implants were printed with hMSCs to assess cell metabolic activity after 1, 4, and 7 days in culture using the alamarBlue assay (Fig. 2H). Fluorescence indicating metabolic activity of cellularized implants was found to increase 5-fold over a 7-day culture period, indicating an increase in cell health, activity, and proliferation.

Additionally, fluorescence imaging of cells (green) and fiber autofluorescence (blue) shows printed implants with a uniform initial distribution of hMSCs throughout after one day in culture and confluent densely-cellularized implants after 26 days in culture (Fig. 2I). Printed cells were found to attach to and begin elongating along the collagen fiber within 24 hours and continued to proliferate to confluency at a rate dependent on cell type, initial cell printing density, and culture conditions. In extended culture, the gross appearance of implants transitioned from largely translucent with visible fiber-like surface texture to an opaque white to yellowish color with a smooth surface texture (Fig. S2), indicating a significant accumulation of deposited ECM. Dense cellular ingrowth as cells bridged gaps between adjacent fibers was also observed, including hMSCs after six weeks of culture on implants sutured into bundles (Fig. 2J) imaged by transmitted light and fluorescence microscopy with cells (green) and fiber autofluorescence (blue).

### Subhead 3: Cell directionality and distribution

The directionality or alignment of cells with respect to a matrix has been shown to affect cellular remodeling potential *(55)* and the alignment of cell-produced collagenous ECM *(56)*. For VML applications, in particular, the alignment of cells within a matrix plays a critical role in facilitating myogenesis and, ultimately, the function of a tissue-engineered muscle construct. As such, directionality analysis in ImageJ was used to quantify cell directionality throughout AC-DC implants using fluorescence imaging and image processing techniques. A representative composite image of a typical 2 mm × 2 mm field of view of an implant printed with MPCs after 14 days of culture is shown in Fig. 3A, with directionality analysis conducted on the fiber-only component of the image (blue) and cell-only component (green) shown in Fig. 3B and Fig. 3C, respectively. Analysis of the fiber-only component shows a narrow frequency distribution, indicating highly parallel fiber with nearly all directional features within ±10° of the peak orientation. Similarly, analysis of the cell-only component shows a frequency distribution with a peak orientation essentially identical to the fiber direction with nearly all components within ±20° of the peak, indicating a significant degree of cell alignment parallel to the fiber. Implementing this analysis across printed implants with both hMSCs and MPCs, we concluded that implants consistently show highly aligned parallel collagen fiber and significant cell elongation in the fiber direction with a high degree of directionality.

**Fig. 3.**
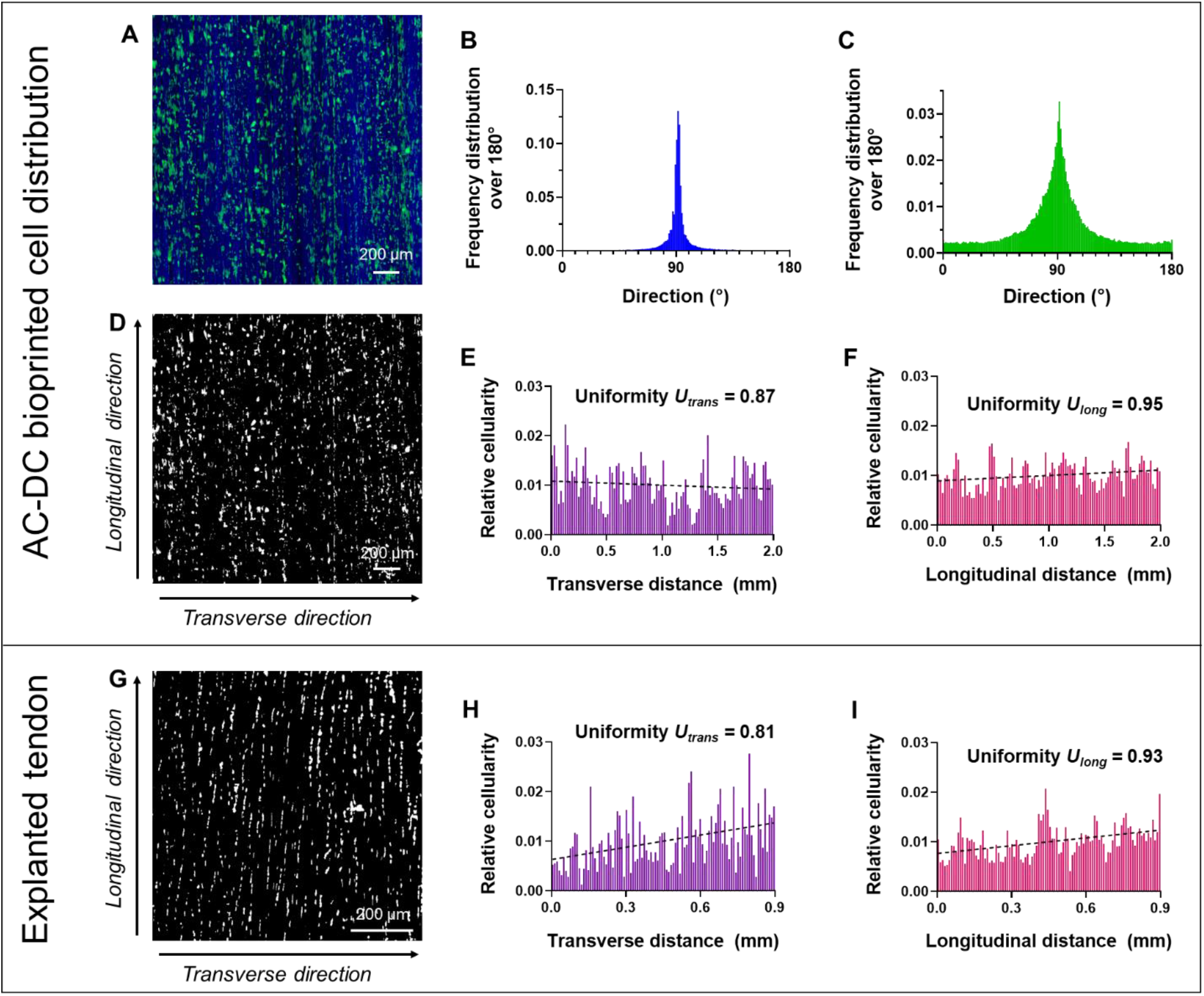
Quantitative cell directionality and distribution. (**A**) Typical field of view of a printed implant with rat muscle progenitor cells (MPCS) distributed throughout and aligned after 14 days in culture. The fluorescence image shows all cells with the cytoplasmic label DiD (green) and collagen fiber autofluorescence at 405 nm (blue). (**B**) Directionality analysis of the fiber-only component of (A), indicating highly parallel fiber. (**C**) Directionality analysis of the cell-only component of (A), indicating a high degree of cell alignment in the fiber direction. (**D**) Image of (A) after processing for cell distribution analysis with cells shown in white, background in black, and transverse and longitudinal directions labeled. (**E**) Relative cellularity plotted along the transverse and (**F**) longitudinal directions, with nearly horizontal linear regression (dashed lines) and quantified uniformity measure *U* (ranging from 0 to 1), indicating a highly uniform distribution of cells. (**G**) Explanted rat Achilles tendon after cryosectioning and fluorescent cell labeling with DAPI, processed for cell distribution analysis with cells indicated in white, background in black, and transverse and longitudinal directions labeled. (**H**) Relative cellularity plotted along the transverse and (**I**) longitudinal directions, with linear regression and quantified uniformity measure *U* indicating a somewhat skewed but largely uniform distribution of cells.

Additionally, we developed methods to quantify the distribution of cells throughout implants by adapting means for analyzing the distribution of particles within a field of view *(57, 58)*. These methods offer a quantitative means to validate AC-DC process control and repeatability and compare and potentially match the designed cellularity of printed implants to native tissue. The results shown herein are representative and illustrate the capabilities of the cell distribution analysis methods. It should be noted that data collected from differing imaging approaches are not intended for direct comparison. Images of printed AC-DC implants with fluorescently labeled MPC cytoplasmic membranes (Fig. 3D) and rat Achilles tendon sections with fluorescently labeled cell nuclei (Fig. 3G) were processed according to our protocols. The relative cellularity, determined by the number of white pixels indicating cellular material compared to black pixels indicating space devoid of cells, is calculated and plotted along the transverse (Fig. 3E, H) and longitudinal directions (Fig. 3F, I) of the images. Plots of relative cellularity offer a means to easily visualize cell distribution throughout printed implants compared to native tissue, with peaks, valleys, and skewness indicating variations in the number and placement of cells throughout a field of view. Linear regression analysis can further be used as a facile method to assess cellularity. For a perfectly uniform cell distribution with data analyzed in 100 bins, linear regression analysis will result in a horizontal line with a *y*-intercept of 0.01. Thus, the relative cellularity of each bin will be one-hundredth of the total number of cells. From a representative field of view of a printed AC-DC implant with MPCs, it is seen that linear regression results in a nearly horizontal line when measured across both the transverse and longitudinal directions (Fig. 3E, F), indicating an essentially uniform distribution of cells throughout the print. Similarly, from a representative tissue section of a rat Achilles tendon, the distribution of cells was found to be somewhat less uniform with a slightly skewed distribution (Fig. 3H, I). Fig. S3 illustrates a case where the distribution of cells is nonuniform and is made apparent by distribution analysis.

We further implemented an additional method to quantify the distribution of cells with a uniformity measure *U* based on Shannon entropy *(57, 58)*. Briefly, the uniformity measure *U* ranges from 0 to 1, where a perfectly nonuniform distribution in which cells are present in exactly one half of a field of view scores a 0, and a perfectly uniform distribution in which cells are present exactly equally throughout scores a 1. As with calculating and plotting relative cellularity, uniformity was calculated across the transverse direction (*U*_*trans*_), and the longitudinal direction (*U*_*long*_) and was determined using the same imaging and image processing techniques. For example, for a representative AC-DC implant with MPCs, cell uniformity analysis yields *U*_*trans*_=0.87 and *U*_*long*_*=*0.95 (Fig. 3D-F). Similarly, from a representative rat Achilles tendon section, analysis yields *U*_*trans*_=0.81 and *U*_*long*_*=*0.93 (Fig. 3G-I), showing high uniformity and even distribution of cells within the bioprint.

### Subhead 4: Implant mechanical properties

We assessed the mechanical properties of AC-DC implants printed with and without hMSCs after 1 day and 28 days in static culture to evaluate the load-bearing capabilities, stability, and effects of cellular remodeling *in vitro*. A custom 2-pin mounting approach for tensile testing (Fig. 4A) was found to provide significantly more consistent results when compared to mounting implants in standard compression grips, which often lead to implant damage, slippage, or staggered breakage of individual fibers within an implant. Representative stress-strain curves for each tested group are shown in Fig. 4B, with distinct “toe” regions of gradually increasing slope followed by linear regions of maximum slope and ultimately well-defined rapid decreases in stress indicating failure. Mechanical properties were calculated and reported using two methods for determining the cross-sectional area of printed implants: “solid only” and “full implant.” Properties determined using the “solid only” cross-sectional area offer theoretical estimates for the constituent collagen fiber itself, based on prior methods *(39, 40)*. Properties determined using the “full implant” cross-sectional area are more representative of an implant’s inherent properties because the void space, limited packing density of constituent fiber, and additional volume due to cell-deposited ECM are considered.

**Fig 4.**
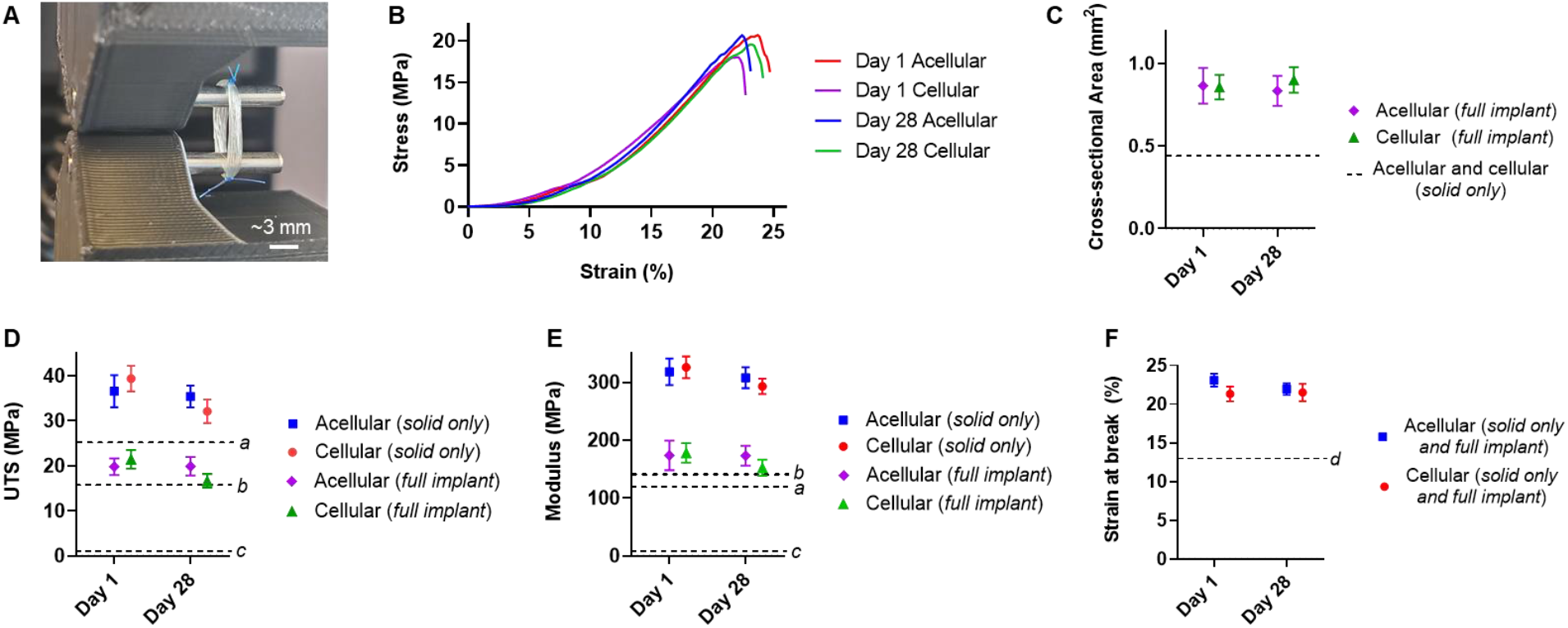
Implant mechanical properties. AC-DC implants were printed with and without hMSCs and assessed after 1 day and 28 days in culture. **(A)** A custom 2-pin uniaxial tensile tensing setup was found to improve consistency in implant failure compared to traditional compression grips. **(B)** Typical stress-strain curves for each experimental group based on “full implant” cross-sectional area. **(C)** Measured “full implant” and “solid only” cross-sectional areas. **(D)** Ultimate tensile stress (UTS). **(E)** Tangent modulus. **(F)** Strain at break. ^*a*^ Mean UTS and modulus of human ACL *(59)*. ^*b*^ Mean UTS and modulus of the strongest portion of the human supraspinatus tendon *(30)*. ^*c*^ Mean UTS and modulus of typical collagen gels used in tissue engineering *(60)*. ^*d*^ Mean peak strain of the human ACL during normal walking *(61)*. (All data n=10 per group per time point).

Cross-sectional area measurements show that the “full implant” area is approximately twice that of the theoretical “solid only” area after both 1 day and 28 days in culture (Fig. 4C). This area measurement is based on the mean hydrated individual fiber diameter of 90.7±6.8 μm and the 72 individual strands of fiber in each implant. These measurements indicate a significant contribution of void space to the full implant cross-sectional area, which can be attributed to the intentionally porous nature of implants where the designed spacing between fibers facilitates nutrient diffusion and prevent shearing or displacement of cells from adjacent fibers. The UTS, tangent modulus, and strain at break were determined using both approaches to determine the cross-sectional area (Fig. 4D-F). For plots displaying UTS (Fig. 4D) and tangent modulus (Fig. 4E), horizontal lines are plotted indicating the mean UTS and tensile modulus of human ACL *(59)*, the strongest portion of the human supraspinatus tendon *(30)*, and typical collagen gels used in tissue engineering *(60)*.

Acellular and cellular implants produced using AC-DC bioprinting nearly match or exceed key mechanical properties of representative native human tendons directly after printing and continue to do so after 28 days in culture. Notably, the UTS and modulus of collagen microfiber implants produced using AC-DC are several orders of magnitude larger than the strength and stiffness of collagen gels commonplace in biomanufacturing approaches, with a typical UTS around 20 kPa and tensile modulus around 200 kPa *(60)*. Both acellular and cellular implants after 1 day and 28 days in culture underwent greater than 20% strain before failure (Fig. 4F), which is independent of the cross-sectional area such that results do not differ between the “solid only” or “full implant” measurement approaches. Thus, AC-DC implants provide sufficient elasticity to withstand typical strain values *in vivo*, such as the peak strain of 13.2% of the ACL during normal walking *(61)*.

### Subhead 5: Functional recovery in a volumetric muscle loss model

*In vivo* skeletal muscle repair studies were conducted over 12 weeks in a validated rodent VML model. At least 20% of overall muscle weight was removed from the tibialis anterior (TA) muscle of the lower left hindlimbs of Lewis rats *(11, 62)*. Three methods of repair were assessed head-to-head: a control group receiving no repair, an acellular implant group receiving repair with AC-DC implants with no cellular component, and a cellular implant group receiving repair with AC-DC implants printed with rodent MPCs. Defect creation, initial placement of an implant, suture placement for implant attachment, and fascia replacement are shown in Fig 5A-D, respectively. All animals recovered post-surgery, and there were no signs of infection and no deaths. Across experimental groups, animal body weight increased similarly over the 12-week period (Fig. 5E), and measured defect weight at the time of surgery was not statistically different (Fig. 5F).

**Fig. 5.**
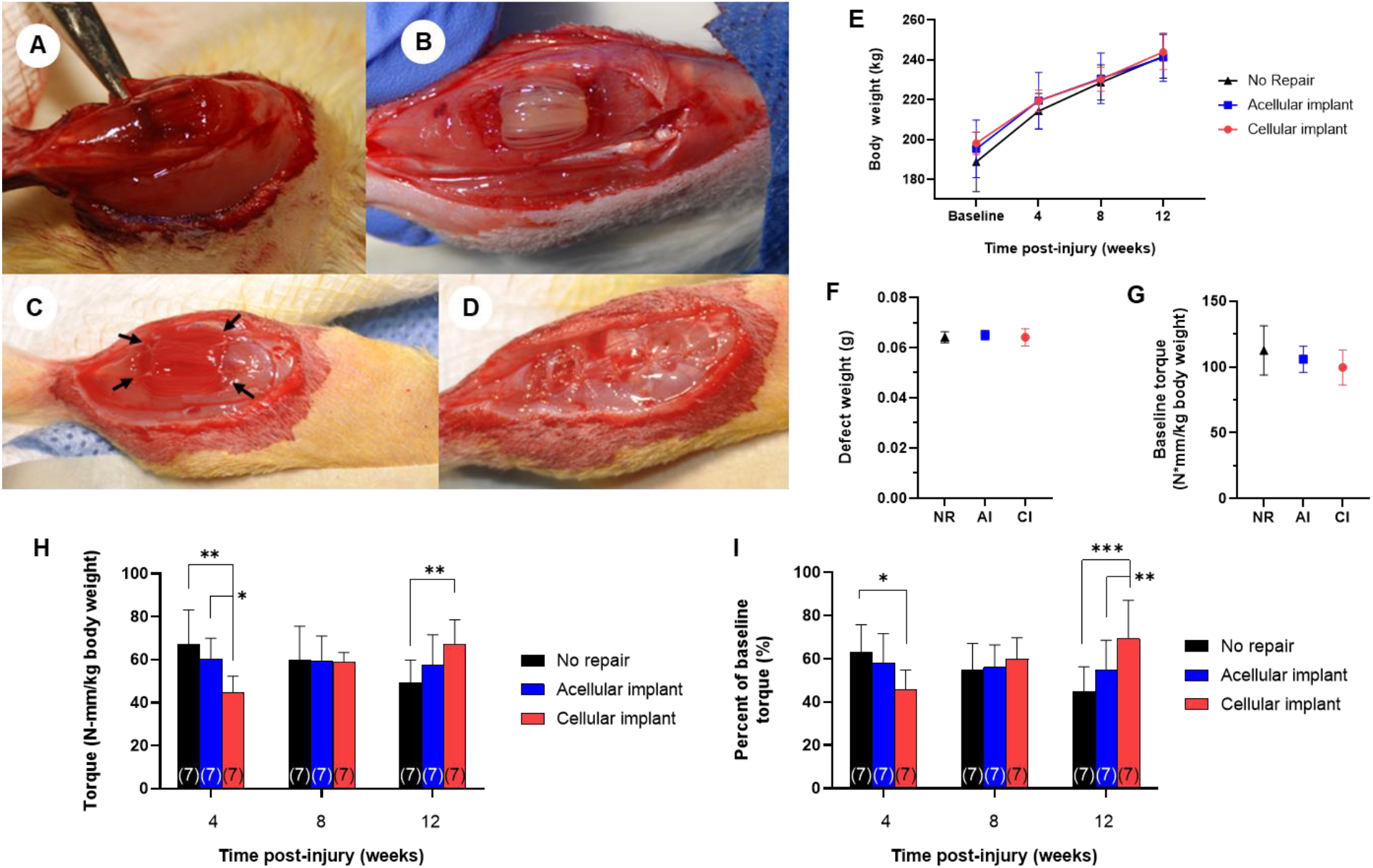
Functional recovery in a rodent VML model. (**A**) Creation of a VML injury measuring approximately 1 cm x 0.7 cm x 0.5 cm and weighing at a minimum 20% of the overall TA weight. (**B**) Acellular AC-DC implant inserted into the injury site and (**C**) sutured into the injury site with arrows indicating attachment points. (**D**) Fascia sutured overtop of the injury site to secure the implant in place further. (**E**) Animal weight pre-injury at 4, 8, and 12-weeks post-injury, corresponding to functional testing timepoints. (**F**) Weight of defects created for No repair, acellular implant, and cellular implant (NR, AI, and CI, respectively) experimental groups (p = 0.8, no significant difference). (**G**) Baseline torque generation pre-injury (p = 0.9, no significant difference). (**H**) Measured torque and (**I**) percent of baseline torque at 4, 8, and 12-weeks post-repair, indicating functional recovery facilitated by implant implantation. (All data n=7 per group per time point, *p<0.05 indicates significance).

Functional testing was performed *in vivo* before defect creation and at 4, 8, and 12-weeks post-repair to assess muscle recovery post-operatively. Briefly, rat hind limbs were attached to a motorized footplate and stimulated electrically to measure maximum isometric torque generation *(11, 54, 62)*. Mean values are expressed as torque normalized to animal body weight at each time point (N-mm/kg of body weight) to control for increases in torque production due to animal growth. Baseline torque generation capability before defect creation did not vary statistically between treatment groups (Fig. 5G). Torque generation post-repair is expressed as raw torque (Fig. 5H) and percent of baseline torque generation (Fig. 5I). Both methods show similar trends with only slight variations in statistical significance.

Most notably, significant improvements in torque generating capability were observed over 12 weeks for injuries repaired with cellularized implants containing therapeutic MPCs. At 4 weeks, raw torque generation was significantly lower in the acellular and cellular implant groups than no repair, and the percent of baseline torque was significantly lower in the cellular implant group. We hypothesize that this initial decrease in torque generation capabilities is due to the early wound healing processes, with possible immunological responses to the introduction of implant materials or cells. However, by 8 weeks post-repair, there was no difference observed between the treatment groups. At 12-weeks post-repair, in contrast to findings at 4 weeks, raw torque generation was found to be significantly higher in the cellular implant group compared to the no repair group, and the percent of baseline torque was significantly higher in both the acellular and cellular implant groups, revealing key trends in the functional recovery of a VML injury among treatment groups. In addition, significant deterioration of function was found over 12 weeks for animals receiving no repair. In contrast, torque generation remained largely consistent for animals repaired with acellular implants, indicating that the presence of the collagen fiber implant without cells attenuated the functional deterioration associated with no repair.

It should be noted that the ablation of synergistic muscles during defect creation removes ∼20% of torque generation in the anterior compartment *(11)*. As such, normalized torque would be limited to ∼85 N-mm/kg across the treatment groups (106 N-mm/kg average at baseline). The mean functional recovery of the cellularized implant group at 12 weeks was 76% of the maximum theoretical recovery following synergist ablation compared to 67% in the acellular group and 57% in the no repair group. In addition, we noted that three of the seven animals receiving repair with cellular implants were observed to have a functional recovery of greater than 87%, with one animal recovering to near-maximal theoretical recovery compared to pre-injury levels (99%).

Following assessment of functional recovery *in vivo* at 12-weeks, isolated TA muscles were collected for morphological and histological examination (Fig. S4). The gross morphology of those repaired by acellular and cellular AC-DC implants appeared more similar to control muscles than did the no repair group, which exhibited convex indentations at the injury location. More fascia was also noted in the repair groups. The distinction between implants and surrounding tissue was not obvious, indicating tissue ingrowth around or resorption of the collagen fiber implants. Isolated muscles were cross-sectioned through the belly and processed for H&E staining, with representative images for each experimental group shown in Fig. 6A-D.

**Fig. 6.**
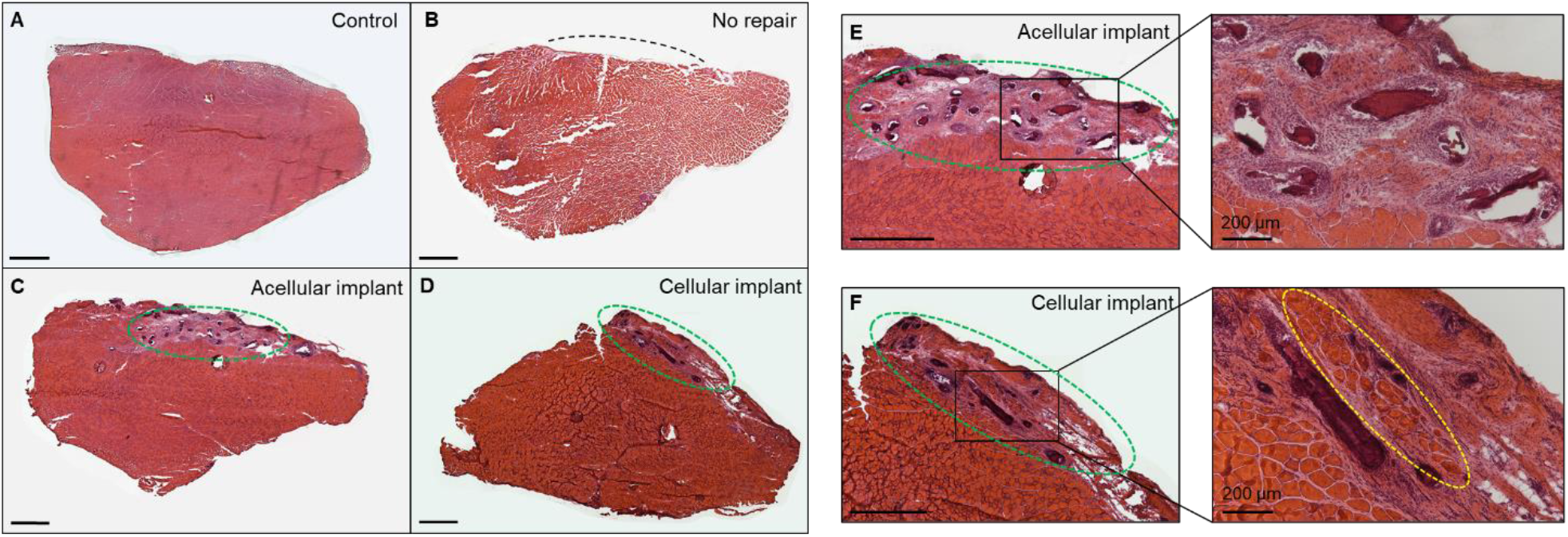
Histological assessment of the TA. Representative H&E images of the TA muscle for (**A**) uninjured control, (**B**) no repair, (**C**) acellular implant, and (**D**) cellular implant experimental groups after 12 weeks. A black dashed line indicates the approximate area of defect creation. Green dashed ovals identify AC-DC implant locations. Magnified views of (**E**) acellular implant and (**F**) cellular implant locations with magnified windowed views showing cellular ingrowth and muscle fiber formation in the cellular implant location (yellow dashed oval). All scale bars are 1 mm unless otherwise noted.

As with gross examination, the unrepaired group exhibited distinct depressions at the injury site indicating a lack of tissue regeneration (Fig. 6B). Animals repaired with acellular and cellular implants, in contrast, exhibited more fullness to the tissue and uniform cross-sections similar to uninjured controls and thus improved cosmesis. Collagen fiber remaining from implants is visible within the injury sites as deep pink somewhat-circular cross-sections on the order of 100 μm diameter. Cellular ingrowth is visible in and around the implants (Fig. 6E and F). Fiber cross-sections are more apparent in the acellular implant group than the cellular implant group, possibly indicating an increased rate of fiber resorption for cellularized implants. For injuries repaired with cellular AC-DC implants, we also note the presence of new muscle fibers at the implant site (Fig. 6F).

Additional sections from the TA muscle belly were processed for analysis using SMASH, a semi-automated muscle fiber analysis software. Laminin and fluorophore 488 staining identify the outline of muscle fibers throughout sections (Fig. 7A-D) and SMASH analysis allows for individual fiber distinction, as seen with colorization applied (Fig. 7E-H). Analysis of the total number of fibers yields no significant difference between the uninjured control, no repair, acellular implant, and cellular implant groups (Fig. 7I). However, the median fiber cross-sectional area (FCSA) in muscle sections repaired with acellular and cellular AC-DC implants was significantly larger than that of the no repair group and did not differ significantly from the uninjured control (Fig. 7J). The cellularized implant and control groups show the greatest difference from the no repair group, with p values of 0.0007 and 0.0002, respectively. Multiplying the total number of fibers by the median fiber cross-sectional area offers a representation of the total muscle fiber cross-sectional area (Fig. 7K). Again, this product shows no significant difference between uninjured controls and injuries repaired with acellular and cellular implants after 12 weeks in life, supporting that AC-DC implants facilitated an increase in total muscle fiber area.

**Fig. 7.**
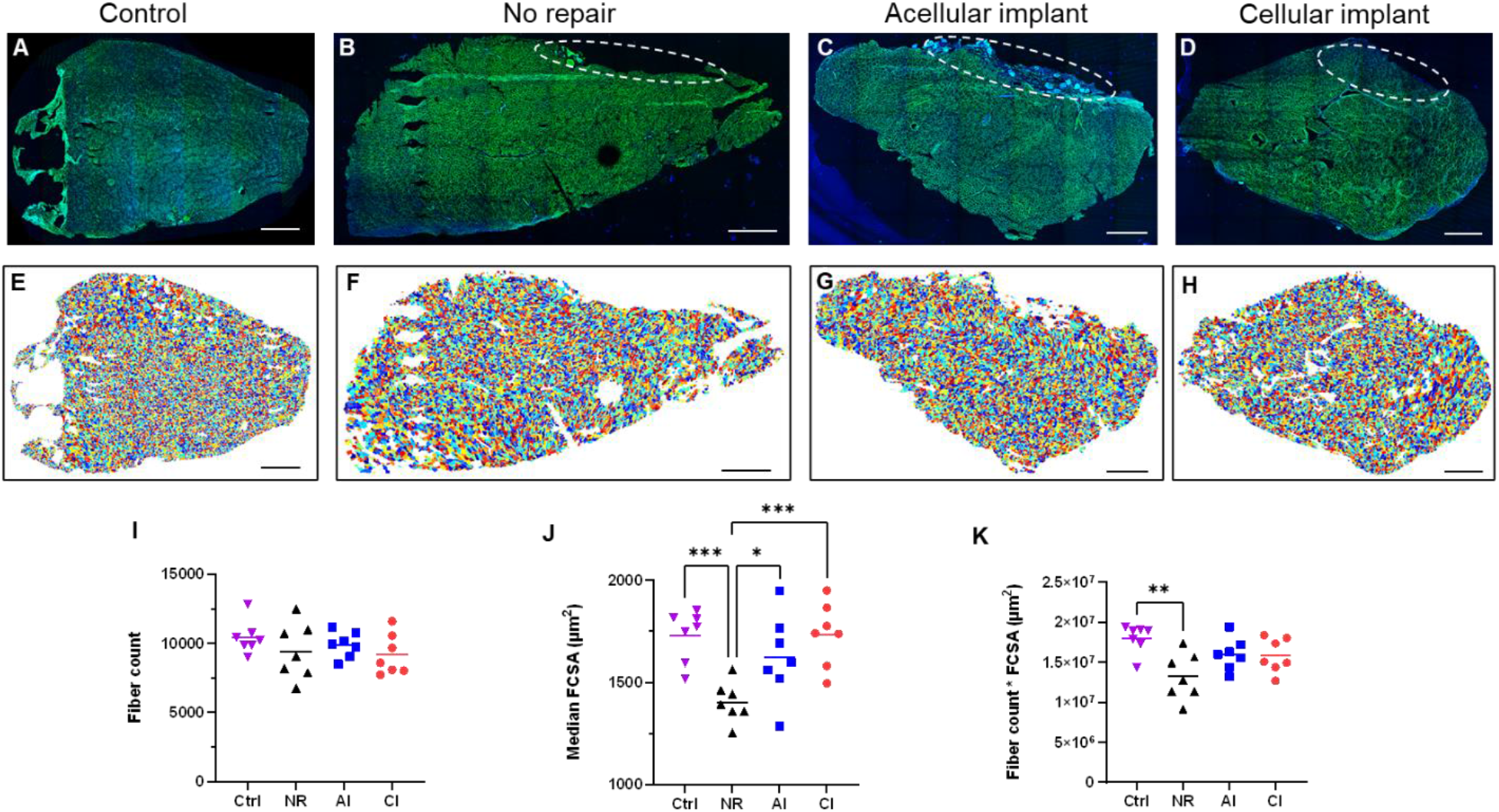
Muscle fiber quantification using SMASH. Representative laminin-stained sections of the TA muscle for (**A**) uninjured control, (**B**) no repair, (**C**) acellular implant, and (**D**) cellular implant experimental groups with dashed ovals indicating the approximate region of injury. (**E**-**H**) Colorized outputs from the software identifying individual muscle fibers within sections corresponding to (A-D), respectively. (**I**) Total fiber count, (**J**) median fiber cross-sectional area (FCSA), and (**K**) the product of fiber count and FCSA for uninured control (Ctrl), no repair (NR), acellular implant (AI), and cellular implant (CI) experimental groups. All scale bars are 1 mm. (All data n=7 per group per time point, *p<0.05 indicates significance).

## DISCUSSION

### Bioprinting concept and design

The AC-DC bioprinting approach rapidly and robustly produced implants consisting of highly aligned collagen microfiber with cells controllably placed throughout. As shown in Fig. 1, the system is built upon a commercial 3D printing platform. It is relatively affordable and straightforward in design, with all modifications completed using off-the-shelf components or parts 3D printed in-house from PLA. While densely packed parallel fiber implants with a rectangular macrostructure were the focus herein, fiber orientation, spacing, and overall implant geometry can be varied using the Python code developed to generate g-code for the AC-DC process. Printed fiber patterns can be designed such that fibers are angled with respect to one another to control material property anisotropy. In contrast, the designed spacing between individual fibers offers a means to control implant porosity. Additionally, fiber can be collected on “frames” of varying geometries rotated by the collection assembly (a cylindrical mandrel, for example), and achievable geometric complexity could feasibly be expanded using a collection assembly capable of multi-axis manipulation. This bioprinting process is particularly well-suited for producing planar, cylindrical, or prismatic geometries. Such geometries are ideal for applications targeting the augmentation or replacement of tendon, ligament, and muscle, where implants of aligned collagen fiber mimic the architecture of these native tissues. For applications in which complex geometries are required, however, AC-DC bioprinting could be augmented with additional substrate geometries to produce more complex shapes and patterns using biotextile or other approaches.

### Implant print fidelity and cellularity

Printing with a single cell suspension, as in Fig. 2, AC-DC implants consist of densely-packed parallel collagen fiber with designed macroscopic dimensions. As Fig. 2 illustrates, the microarchitecture and topography of AC-DC implants were shown to facilitate and guide cell attachment, alignment, and proliferation *in vitro*. We envision that this microstructural guidance for both printed and endogenous cells in the case of implantation will ultimately lead to improved tissue strength for ligament and tendon applications and functional contractility for VML applications. Printed implants exhibited a high degree of directionality for collagen fiber and cells analogous to native musculoskeletal tissues (Fig. 3A-C). In addition to impacting cellular remodeling potential *(55)* and cell-produced ECM organization *(56)*, cell and matrix alignment may play a critical role in organized tissue biointegration and cell differentiation. For hMSCs in particular, an aligned collagen matrix has been shown to significantly upregulate the expression of tendon-specific markers, including scleraxis, tenomodulin, tenascin-C, and collagen-III compared to a randomly oriented matrix *(63)*.

### Cell directionality and distribution

Qualitative analysis methods were implemented to assess both cell orientation and distribution throughout the implants (Fig. 3). The validity and significance of both techniques are reliant on consistent use of fluorescence labeling, image collection, and image processing techniques. The qualitative analysis shown herein enables the characterization of printed implants and the standardization and quality assessment of the AC-DC bioprinting process. These methods may also be applied to various tissue engineering approaches traditionally relying on subjective and qualitative descriptions.

Cell labeling plays a significant role in the interpretation of quantitative cell orientation and distribution analysis and is a methodology lacking standardization in the bioprinting field that is helpful to assess print quality. Our approach provides an unbiased and quantitative means of accessing bioprint fidelity in a novel way to address this deficiency in the field. Our described systems quantify our AC-DC bioprints (Fig. 3) with uniformity measure *U* nearing 1 found for both representative cases, indicating a largely uniform and homogeneous distribution of cells when analyzed across both the transverse and longitudinal directions. Alongside calculation of relative cellularity, these analysis methods provide quantitative measures describing the placement and overall homogeneity of cells throughout bioprinted implants or sections of native tissue. These measures offer a powerful means of comparing batch-to-batch variation in fabricated constructs and comparing the spatial cellular properties of native tissue targeted for repair or augmentation, and to compare results among products, bioprinters, and labs. While the elongation of cell nuclei can be observed in aligned cells, techniques that enable visualization of cell body orientation such as cytoplasmic or cytoskeletal labeling provide significantly improved results. However, cell distribution analysis can be conducted with cytoplasmic labeling (Fig. 3D-F) or cell nuclei labeling (Fig. 3G-I). Generally, cytoplasmic labeling enables analysis at a lower magnification and over a larger field of view. In contrast, cell nuclei labeling offers improved accuracy due to minor cell-to-cell variation in the size of the nucleus compared to the cytoplasm. Cell orientation was also found to impact cell distribution analysis when using cytoplasmic labeling. More severe peaks and valleys are present in the measured relative cellularity across the transverse direction compared to those measured across the longitudinal direction (Fig. 3D-F). Changes in cellular orientation can be attributed to cell elongation, which acts to increase the measured grayscale value in the direction of elongation (pixel columns measured across the transverse direction). Regardless, both labeling approaches were found to enable meaningful quantification of cell distribution throughout a field of interest, whether a printed implant or a native tissue section.

### Implant mechanical properties

As shown in Fig. 4, critical mechanical properties of AC-DC implants both with and without cells and after 1 and 28 days of culture were found to closely match or exceed properties of the human ACL *(59)* and supraspinatus tendon of the rotator cuff *(30)*, and offered properties ∼1000 times those of typical collagen gels *(60)* used in tissue engineering and bioprinting. Interestingly, the stress-strain curves of all experimental groups exhibited a distinct “toe” region, indicating a gradual increase in stress in response to tensile loading, which is often observed when testing explanted tendons and ligaments. While this behavior is attributed to the un-crimping of collagen fibrils in native tissue, herein, it is likely due to the non-simultaneous recruitment of constituent fiber throughout printed implants. Although by a different mechanism, the presence of a toe region may indicate that AC-DC implants further mimic the functional behavior of native tissues to act as biological “shock absorbers” to suddenly applied loads.

Perhaps most notably, AC-DC implants offer the structural and mechanical properties required to withstand and transmit significant loads in the body and feasibly act as clinically relevant tissue augmentations immediately after printing. This progresses beyond current biomanufacturing approaches, which often require months of complex and costly bioreactor culture to improve the mechanical properties of produced constructs *(31–33)*. Due to the extremely high initial strength of AC-DC implants, we have not observed an increase in mechanical properties over 28 days of static culture and observed slight but statistically significant decreases in the strength and stiffness of cellularized implants. While acellular implants did not experience reductions in properties over 28 days, indicating the *in vitro* stability of the collagen microfiber alone, we hypothesize that the decreases in cellular implant properties can be attributed to cell-induced degradation and remodeling of the collagen fiber. From gross observation, cells were found to have bridged gaps between adjacent fibers and created a more opaque, smooth implant appearance as in Fig. 2J. In addition, while fibers were bound to one another by cells and newly synthesized ECM, we did not observe an increase in tensile strength or stiffness. Thus, we theorize that cell-induced degradation of the collagen fiber has outweighed any potential increase due to newly deposited ECM, primarily due to the difficulty of ECM materials matching the extremely high initial strength and stiffness of the collagen microfiber.

### Functional recovery in a volumetric muscle loss model

*In vivo* assessment of AC-DC implants revealed a substantial capacity for recovery in a volumetric muscle loss injury model, with acellular implants preventing functional deterioration associated with no repair and, most notably, cellular implants with MPCs facilitating a significant increase in torque generating ability (Fig. 5). Histological and computational analyses support that the mechanism for improved functional regeneration over 12 weeks is the formation of new muscle fibers (Fig. 6F) and increase in total muscle size (Fig. 7K). While all experimental groups showed a significant number of new cells in the area of the injury, the group receiving no repair did not exhibit the same maturity of cells as in groups receiving AC-DC implants. This was supported by the SMASH analysis, with the total fiber count between groups found to be similar (Fig. 7I) but the median fiber cross-sectional area in groups receiving implants found to be substantially higher (Fig. 7J). While implants with mechanical properties approximating those of tendon and ligament tissues were effective herein, performance may potentially be further improved by producing implants with properties more closely matching those of muscle.

### Interpretation of results and clinical translation

We have developed AC-DC bioprinting to produce regenerative collagen fiber-based implants that can be incorporated into the surgical management of a number of musculoskeletal injuries to restore lost soft tissue structure and function. In particular, tears at the myotendinous junction of the muscle-tendon unit as well as en bloc muscle defects have remained vexing injuries with few, durable reconstructive solutions. Production and analyses were done at a scale appropriate for rodent-sized implants, yet processes can readily be scaled up to produce human-scale implants. In conjunction with scale-up, techniques to facilitate implant handling and surgical delivery are needed for many surgical implants. For prospective human clinical use, critical quality attributes of AC-DC implants are important next steps. The *in vitro* analysis methods implemented herein provide valuable tools for quantifying cellular and mechanical properties, useful for future validation work. While acellular implants showed some efficacy in the VML model, only bioprinted implants containing therapeutic cells facilitated a significant degree of functional recovery. Although there are complexities associated with clinically translating cell-based biomanufactured products, AC-DC bioprinting has the potential to become a platform technology capable of producing implants with revolutionary regenerative capabilities to address a range of musculoskeletal and other disorders.

## MATERIALS AND METHODS

### Subhead 1: Study design and reproducibility

#### Rodent VML model

Choice of sample size (n=7 animals per group per time point) was informed by our previous studies and experience with regenerative implants in a rat TA injury model *(11, 62, 64)*. This study was intended to provide robust statistics to determine significant differences between groups for quantitative *in vivo* functional analysis. Isometric contractile torque has historically resulted in the greatest variability among treatment groups and is the determining factor for the required number of animals. Power analysis assuming a 30% difference (standard deviation 20%) between treatment and control groups indicated that 7 animals per group achieves 80% power at alpha = 0.05. The number of animals was not altered over the course of the study and endpoints were determined before study initiation. No animals exhibited signs of pain and distress or required euthanasia before designated study end points. Treatment groups were randomized and quantitative analysis methods were standardized to minimize any undue bias.

### Subhead 2: Experimental methods

#### Collagen fiber production and sterilization

Collagen microfibers for use in AC-DC bioprinting were produced using a wet-spinning microfluidic extrusion process, as described in detail in our previous work *(43)*. Briefly, clinical-grade lyophilized collagen was dissolved in acetic acid. Next, acidified collagen was extruded through microneedles into a bath of a neutralizing alkaline formation phosphate buffer containing salts and polyethylene glycol and passed through a bath of aqueous ethanol. The fiber was collected on spools for subsequent glyoxal crosslinking, sterilization by electron beam sterilization (Steri-Tek, Fremont, CA) using a 20±2 kGy target dose, and stored desiccated until use.

#### hMSC and related cellularized implant culture

Human bone marrow-derived mesenchymal stem cells (hMSCs) (RoosterBio^®^, Ballenger Creek, MD) and cellularized implants produced using hMSCs were cultured and maintained in serum-free, xeno-free media (RoosterBio^®^). Cell culture media was supplemented with 1-2% Gibco™ Antibiotic-Antimycotic (ABAM) (Thermo Fisher Scientific). Cells and implants were prepared and handled aseptically and cultured maintaining physiological conditions at 37°C and 5% CO_2_. Implants remained fully submerged during culture, and two-thirds of the total media volume was replaced every 2-3 days. For comparative studies, acellular implants were cultured under conditions identical to those of cellularized implants.

#### MPC and related cellularized implant culture

Muscle progenitor cells (MPCs) were isolated as previously described *(62)*. The tibialis anterior (TA) and soleus muscles of 4 weeks old female Lewis rats (Charles River Laboratories, Wilmington, MA) were excised, sterilized in iodine, and rinsed with PBS (Hyclone). Muscles were minced to create a homogenous cell slurry and then incubated for 2 h at 37C in 0.2% collagenase (Worthington Biochemicals, Lakewood, NJ) in DMEM. The homogenous muscle slurry was then pre-plated onto 10-cm collagen-coated tissue culture dishes (Corning, Corning, NY) at 37C in a myogenic medium containing DMEM high glucose with 20% FBS, 10% horse serum (Gibco by Life Technologies); 1% chick embryo extract (Accurate, Westbury, NY), and 1% ABAM. After 24 h, the cell suspension was transferred to 15-cm Matrigel-coated (1:50; BD Biosciences, Franklin Lakes, NJ) tissue culture dishes. Cells were passaged at 70–90% confluence and further cultured in proliferation media containing DMEM low glucose with 15% FBS and 1% AA. At passage 1, MPCs were frozen using proliferation media supplemented with 5% DMSO and subsequently shipped to Embody, Inc. for further passaging. AC-DC implants printed with MPCs were cultured in myogenic media as described above. Acellular and cellular implants were printed 2-3 days before transportation to University of Virginia by car in a warmed and vented container (approximately 3-hour travel time), cultured 2-3 days before implantation. Cells and implants were prepared and handled aseptically and cultured maintaining physiological conditions at 37°C and 5% CO_2_. Implants remained fully submerged during culture, and two-thirds of the total media volume was replaced every 2-3 days. For comparative studies, acellular implants were cultured under conditions identical to those of cellularized implants.

#### Preparation of cell suspensions for AC-DC bioprinting

Hyaluronic acid (HA) sodium salt from Streptococcus equi (Sigma-Aldrich, St. Louis, MO) was dissolved in appropriate cell type-dependent culture media to a final concentration of 5 mg/mL in a conical glass vial gently stirred overnight. Immediately before printing, cells were harvested and resuspended within the hyaluronic acid solution to desired concentrations by gentle pipetting. Cell suspensions remained at room temperature during printing. Typical printed concentrations ranged from 1×10^6^ cells/mL to 5×10^6^ cells/mL.

#### AC-DC process control and commercial printer customization

Collagen fiber is fed through the seeding manifold and attached at an initial anchoring point on the collection assembly to initiate bioprinting. As the collection assembly motors rotate the frame, fiber is drawn under tension through the seeding manifold and coated by the extruded cell suspension. By coordinating the rotation of the frame and linear translation of the printhead along the width of the frame (the feed), cellularized implants of dense, highly aligned collagen microfiber are produced. A custom Python code was developed to accept user inputs for designed implant geometry and printing parameters, including the number of implants per frame, implant width, length, number of layers of fiber, the extruded volume of cell suspension per millimeter drawn fiber, feed distance between parallel fibers, and frame rotation rate, and output a corresponding g-code file. The g-code file contains all parameters and motion/extrusion commands to execute a designed print and is sent to the printer to produce the designed implant. Repetier-Host is used as a user interface to execute these commands and manual homing, motion, and extrusion commands.

Folger Tech FT-5 R2 commercial 3D printer hardware and firmware were highly modified to facilitate the AC-DC biofabrication approach. The fused deposition modeling printhead was removed and replaced with custom printhead components (Fig. 1A). Non-stock components for the printhead and collection assembly were 3D printed in-house from PLA or machined. All stepper motors and drive pulleys were replaced to improve the resolution on *X, Y*, and *Z* axes. The printer *Z-axis* control was repurposed as a new *R* rotational axis for the custom collection assembly and mounted to the printer’s build plate. The printer firmware was modified accordingly to accommodate hardware and control scheme changes. All components of the AC-DC printing approach contacting implants during printing were autoclaved or soaked in 100% ethanol for at least one hour. The printing and auxiliary handling processes were conducted in either a biosafety cabinet or HEPA filtered laminar flow hood to maintain aseptic environments.

#### Fluorescent labeling, microscopic imaging, and cell viability

According to recommended manufacturer protocols, the fluorescent label Invitrogen Molecular Probes Vybrant™ DiD (Thermo Fisher Scientific, Waltham, MA) was used according to recommended manufacturer protocols to visualize cell cytoplasmic membranes. According to recommended manufacturer protocols, the fluorescent label Ethidium Homodimer-1 (EthD-1) (Thermo Fisher Scientific, Waltham, MA) was used according to recommended manufacturer protocols to visualize dead cell nuclei. Bright autofluorescence of the collagen fiber at 405 nm allowed for clear visualization of fiber throughout implants. All fluorescence and transmitted light microscopic imaging was conducted on an inverted light microscope (Axio Vert.A1 Model, Zeiss, Germany), and composite images were produced using ZEN imaging software (Zeiss, Germany). The quantitative viability of hMSCs in AC-DC implants was assessed immediately after printing, where the total number of cells in a field of view was determined by DiD labeling. The number of dead cells was determined by EthD-1 labeling. Representative fields of view from the central region of each implant were taken for live/dead counting, with results presented as mean ± standard deviation (n=6).

#### Metabolic activity assay

The alamarBlue® assay (Bio-Rad, Hercules, CA) was used to assess the metabolic activity of AC-DC implants printed with hMSCs according to recommended manufacturer protocols. After 1, 4, and 7 days of culture, implants were incubated in an alamarBlue working solution for 4 hours, and sample media and blanks were collected in triplicate. Media was analyzed for fluorescence intensity at 560/590 nm excitation/emission using a Synergy HTX multi-mode microplate reader.

#### Quantifying cell distribution and uniformity

Appropriate fluorescence labeling techniques were applied to visualize cells throughout printed implants, and tissue sections and images were converted to binary to visualize cells in black on a white background. The “Plot Profile” function of ImageJ FIJI (NIH Shareware, Bethesda, MD) using was then used to measure and plot the average grayscale value, with black pixels measured as 255 and white pixels measured as 0, of pixel columns taken across the transverse direction of the field of view, with the value for each column being the average grayscale value of pixels contained in that column. Similarly, the grayscale value is measured for pixel rows taken along the longitudinal direction. The value for each row is the average grayscale value of pixels contained in that row. A custom Python code was developed to group measured greyscale values into 100 bins and normalized to the total grayscale value of the field of view to represent the relative cellularity or cell material found in each transverse or longitudinal bin relative to the entire field of view.

Based on prior applications *(57, 58), U* (Equation 1) defines the uniformity measure based on the Shannon entropy of a printed implant, *s*_*printed*_, Shannon entropy of a perfectly nonuniform particle distribution, *s*_*nonuniform*_, and Shannon entropy of a perfectly uniform particle distribution, *s*_*uniform*_. A perfectly nonuniform distribution is one in which particles entirely fill exactly half of a field of view. A perfectly uniform distribution is one in which particles are present equally throughout. Shannon entropy *s* (Equation 2) is calculated for a field of view with *N* pixel columns/rows across a field of view based on the probability *p*_*j*_ (Equation 3) that a particle lies within the *j*th pixel column/row, where *m*_*j*_ is the grayscale value of the *j*th column/row and *M* is the sum of grayscale values across the field of view. Thus, the uniformity value *U* ranges from 0 for a perfectly nonuniform distribution in which cells are present in exactly half of the field of view to 1 where cells are present exactly equally throughout the entire area. A custom Python code was developed to calculate *U* for a given field of view.

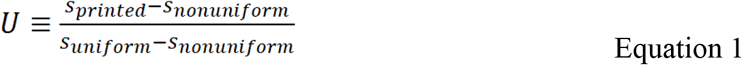

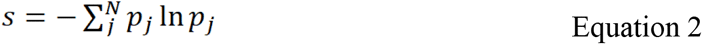

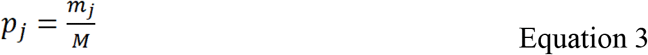

#### Mechanical property testing

Implants were secured into bundles with a suture at each end, mounted to a custom 2-pin mechanical testing setup (Fig. 4A), and pulled to failure under uniaxial tension using a uniaxial tensile testing machine (MTS Systems Corporation, Eden Prairie, MN) with a 100 N load cell. Implants were tested after 1 day and 28 days of culture. Pins were displaced at a rate of 1 mm/s, and load-displacement data were recorded continuously. The “solid only” theoretical cross-sectional area was determined by multiplying the average cross-sectional area of individual collagen fibers measured using microscopic imaging by the total number of constituent fibers within the printed implants *(39, 40)*. “Solid only” cross-sectional area was assumed to be identical for each experimental group based on the consistency and stability of the collagen fiber. “Full implant” cross-sectional area was measured using digital calipers for each sample immediately before testing. Ultimate tensile stress (UTS) was determined by the peak load and cross-sectional implant area, and tangent modulus or stiffness was determined by the slope of the linear region of the stress-strain curve. The strain was determined from the linear pin displacement during testing, and the gauge length was taken as the initial center-to-center distance between the two pins, with strain at break measured at the peak recorded load. A sample number of n=10 was wound to offer statistical significance and consistency based on prior experiments.

#### VML injury model

In vivo skeletal muscle repair studies were conducted over 12 weeks using a total of 21 female Lewis rats (11–14 weeks old with a mean bodyweight of 198.4 g – 3.5 g) (Charles River Laboratories) split into three groups. A VML defect was surgically created in the TA muscle of each rat as previously described *(11, 54, 62)*. A longitudinal incision was made along the lateral side of the left lower hindlimb, from the ankle to the knee. The skin was separated by blunt dissection from the underlying fascia. The fascia was cut and separated from the anterior crural muscles. The extensor digitorum longus (EDL) and extensor hallicus longus (EHL), synergist muscles in the anterior compartment, were isolated and ablated at the tendons to assess the impact of VML injury to the TA. The VML injury was created by excising the middle of the TA in an area measuring approximately 1 cm x 0.5 cm x 0.7 cm and avoiding the underlying tendon. The defect size was calculated as 20% of the TA, which was determined experimentally to be 0.17% of the animal’s body weight. Immediately post VML injury, each group (n=7) received a different treatment, with the control group receiving no treatment, the acellular implant group receiving a collagen-based implant with no cellular component, and the cellularized implant group receiving complete regenerative constructs incorporating muscle progenitor cells. Implants were bound and sutured into the wound bed at the corners of the wound boundary region. (6-0 Vicryl; Ethicon, Somerville, NJ). The fascia was also closed with 6-0 vicryl interrupted sutures. In addition, the skin was closed with 5-0 prolene (Ethicon) interrupted sutures and skin glue to prevent reopening of the incision. Buprenorphine (0.05 mg/kg) was administered subcutaneously for 3 days post-surgery. All animals in this study were treated per the Animal Welfare Act and the Guide for the Care and Use of Laboratory Animals. All procedures were approved by the University of Virginia Animal Care and Use Committee.

#### In vivo functional testing

Functional testing was performed on all animals prior to VML surgery to establish baseline torque responses for each animal. Testing was again conducted post-healing at three time points of 4, 8, and 12 weeks post-surgery. *In vivo*, functional analysis was performed to assess recovery post-VML injury. Torque production of the subject’s TA was measured in vivo as previously described *(11, 54, 62)*. Rats were anesthetized (2% isoflurane; Henry Schein, Dublin, OH), and the left hindlimb was aseptically prepared. The rat was placed on a heated platform, and the left foot was secured at a 90 angle to a footplate attached to the Aurora Scientific 305C-LR-FP servomotor (Aurora, ON, Canada). Two sterilized percutaneous needle electrodes (Chalgren, Gilroy, CA) were inserted superficially through the skin to stimulate the left peroneal nerve. An electrical stimulus was applied (Aurora Scientific Stimulator Model 701C), and stimulation voltage and electrode placement were optimized with continuous 1 Hz twitch contractions. Contraction of the anterior crural muscles leading to dorsiflexion of the foot was determined by measuring the maximal isometric tetanic torque over a range of stimulation frequencies sufficient to result in the plateau of the torque response (10– 150 Hz). At 12 weeks, animals were euthanized through CO2 inhalation and injured TA muscles, and contralateral control muscles were explanted, weighed, and frozen in preparation for imaging.

#### Histological and immunofluorescence examination

TA muscles were explanted at sacrifice. All tissues were flash-frozen in liquid nitrogen and stored at -80C until ready for sectioning. A cryotome was used to create 12 mm cross-sections of the experimental and control TA muscles for all animals in each group at 12 weeks. Hematoxylin and Eosin and Masson’s Trichrome stains were performed following standard procedures to assess cellular morphology and fibrosis, respectively. Images were captured at 4x and 10x (Nikon Upright Microscope). Histological images were examined, particularly in the region of injury, to identify new fiber formation, cellular proliferation, growth, or potential fibrosis.

Other cross-sectional images were stained with immunofluorescent fibrin antibody stain conjugated to Alexa Fluor 488. All antibodies were diluted in Dako Antibody Diluent (Dako Antibody Diluent S0809; Agilent Technologies). Images were captured by confocal microscopy (Leica DMi8; Buffalo Grove, IL). These images were enhanced and uniformly edited using the CLAHE (contrast limited adaptive histogram equalization) plugin of ImageJ (NIH). Analysis was conducted to determine the fiber cross-sectional area and the minimum Feret diameter. These area measurements were accomplished using the software SMASH, a semi-automatic muscle analysis program built in MATLAB (MathWorks), developed at the University of Pennsylvania. The SMASH software, its functions, and limitations have been thoroughly detailed *(65)*. In the 12-week samples that were analyzed in this study, the site of the original VML defect was located by the presence of fibrosis on the surface of the muscle, and in all cases, either an apparent concave depression on the surface of the muscle or detectable, albeit sometimes modest disruptions of the usually smooth surface architecture of the surface of the TA muscle. The distinct anatomical structure of the TA due to its close association with the tibia also allowed for ease in determining where the original defect was made.

#### Subhead 3: Statistical analysis

All data are presented as mean ± standard deviation unless otherwise noted. Sample group sizes are given in respective figure captions, main text, or Materials and Methods sections. Where appropriate, the statistical significance of functional recovery data was determined by one-way ANOVA or two-way ANOVA with multiple comparisons and Fisher’s LSD. SMASH data was analyzed using the Brown-Forsythe and Welch ANOVA test with multiple comparisons.

## Supplementary Materials

**Fig. S1.**
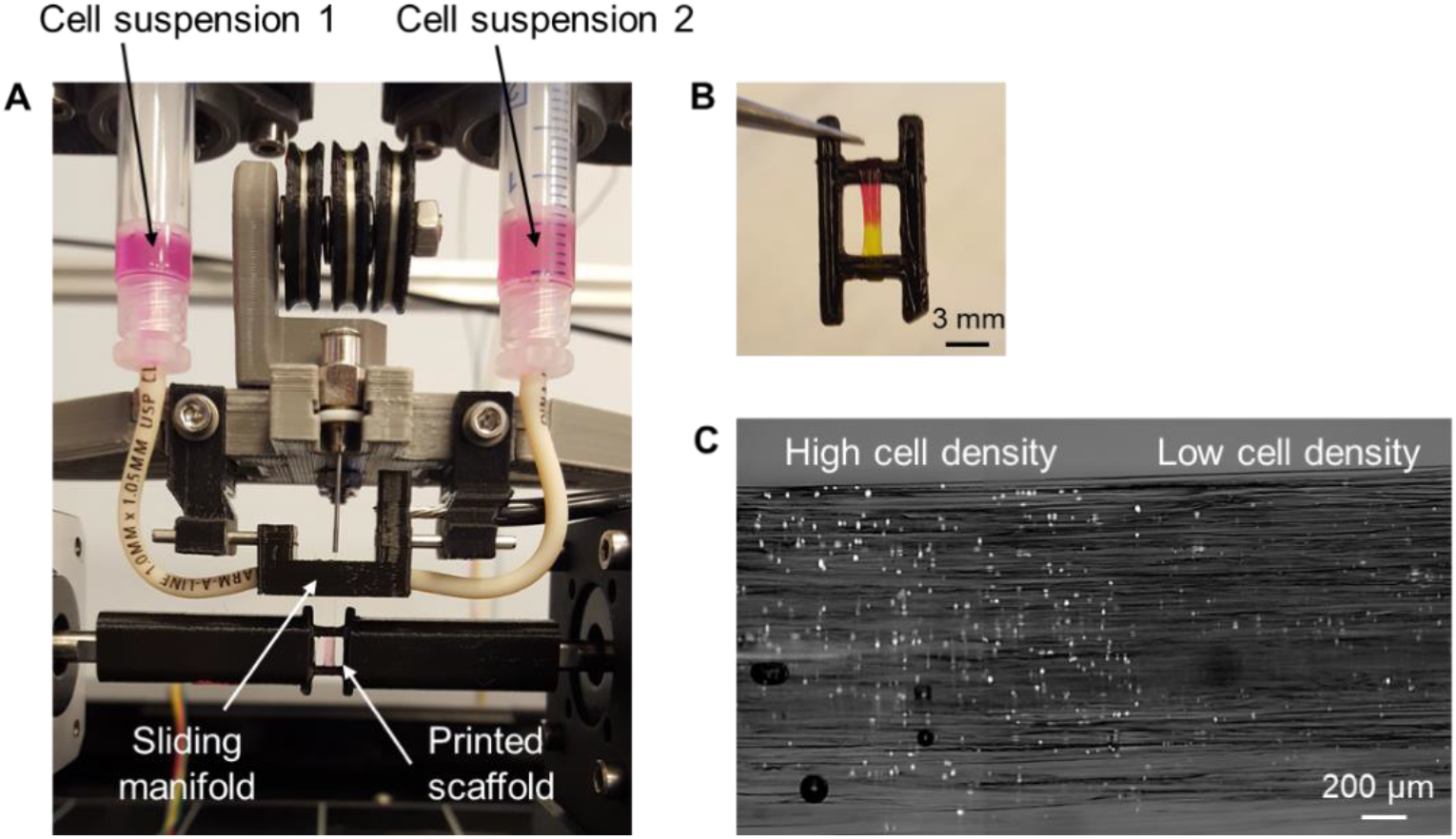
Dual-solution AC-DC proof of concept. (**A**) Dual-solution printhead for AC-DC to produce heterogeneous implants with cells from two distinct cell suspensions. (**B**) Proof of concept implant printed with one pink dyed and a second yellow dyed solution, illustrating a clear distinction between printed regions. (**C**) Fluorescence image of an implant printed using two suspensions of hMSCs, with a high cell density suspension used to print cells in one region (left) and a low cell density suspension used to print cells in the adjacent region (right). Cells were stained with the fluorescent label DiD and imaged immediately after printing.

**Fig. S2.**
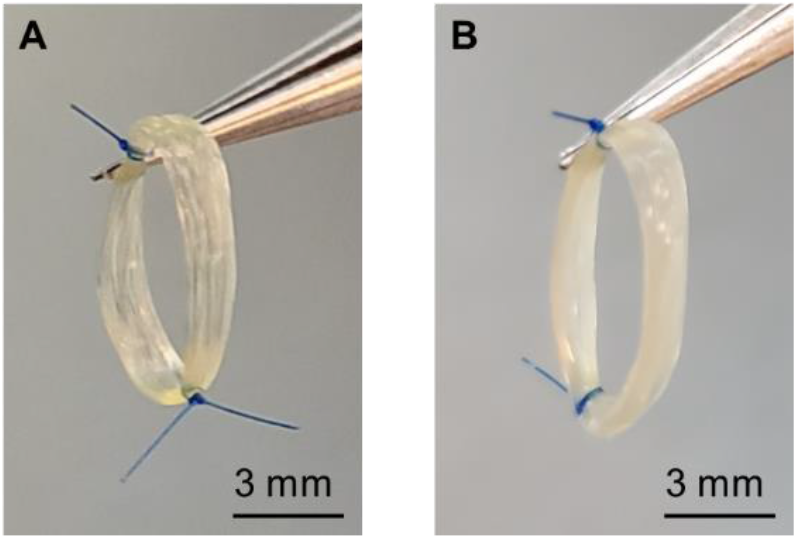
The difference in the appearance of acellular and cellular implants. (**A**) Tweezers hold a typical acellular implant secured by suture immediately after printing. (**B**) A typical raft printed with hMSCs after 14 days of culture held by tweezers, illustrating a smooth and opaque appearance due to cellular material and deposited ECM.

**Fig. S3.**
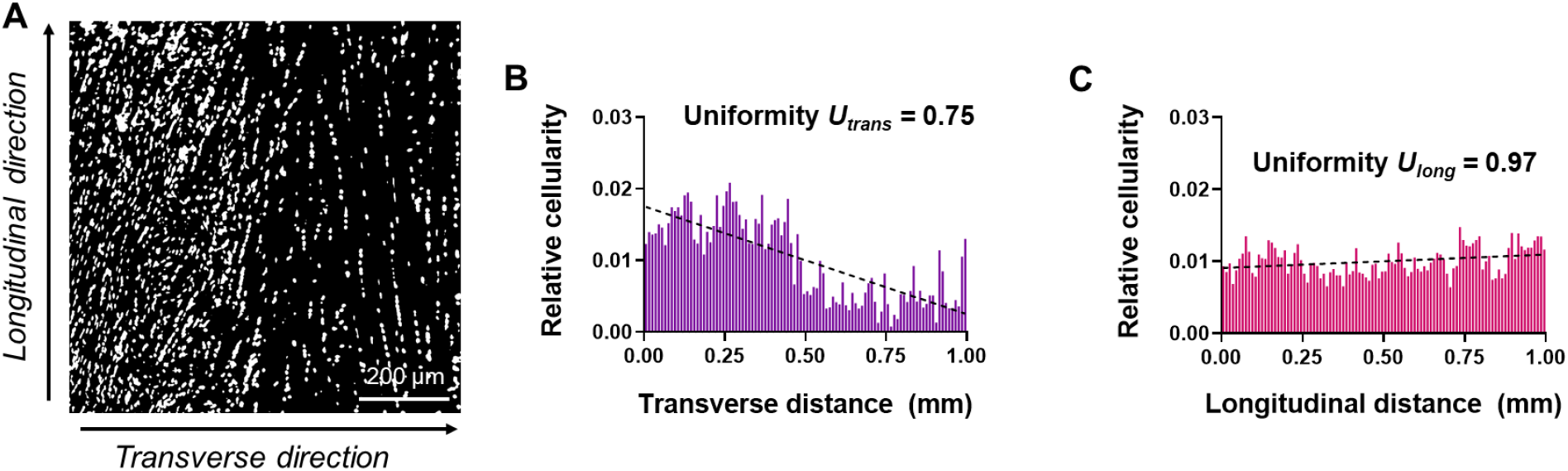
Relative cellularity and uniformity of tissue interface. (**A**) Explanted rat Achilles tendon after cryosectioning and fluorescent cell labeling with DAPI, processed for cell distribution analysis with cells indicated in white, background in black, and transverse and longitudinal directions labeled. The section includes the interface between high and low cell density regions. (**B**) Relative cellularity plotted along the transverse and (**C**) longitudinal directions, with linear regression and quantified uniformity measure *U*. The relative cellularity plot in (B) captures the distinct regions of high and low cell density.

**Fig. S4.**
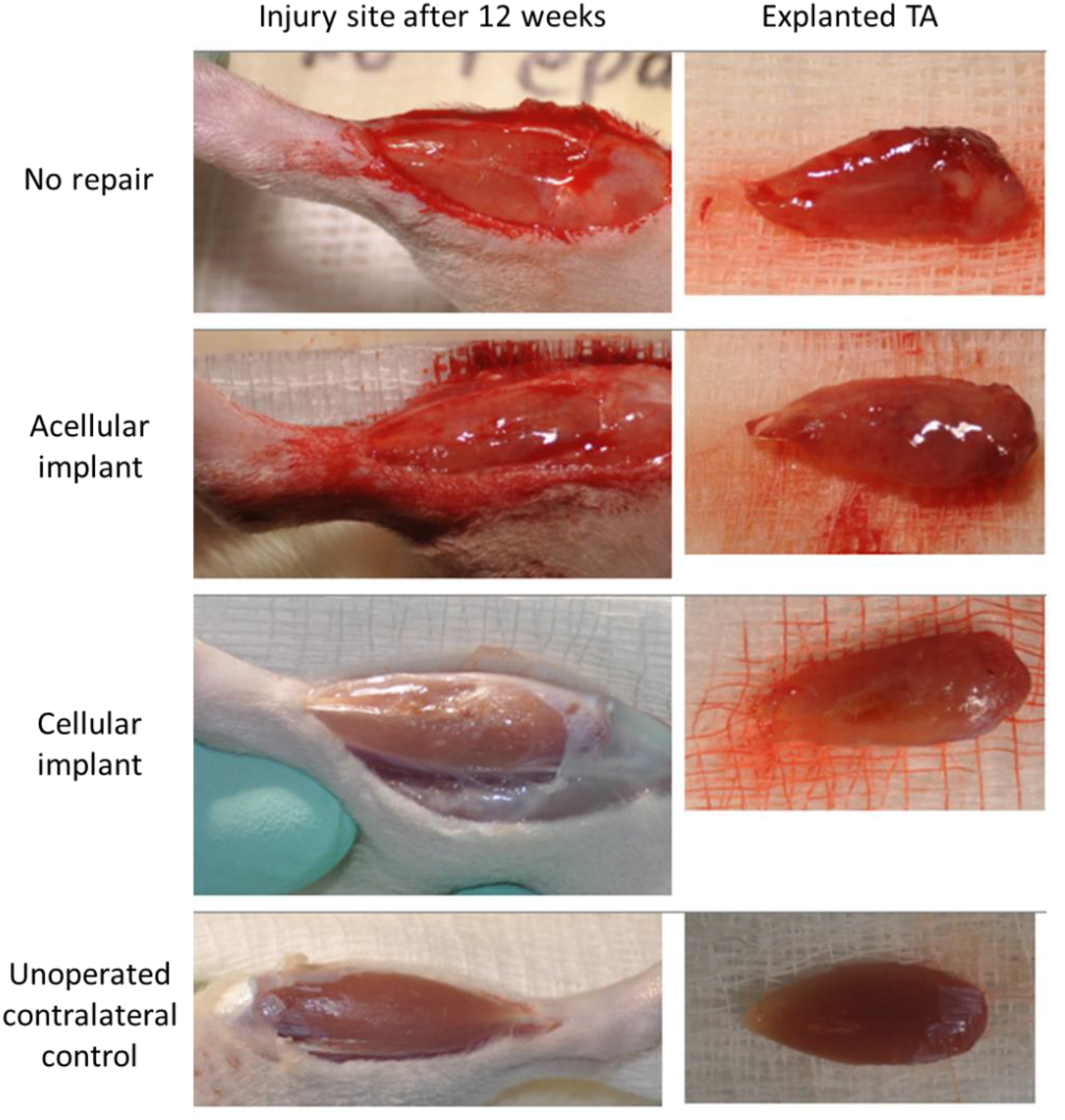
Gross morphology of tibialis anterior muscles 12 weeks post-repair. Exposed tibialis anterior (TA) muscles of each experimental group before (left) and after (right) being explanted for further assessment.

**Movie S1.**
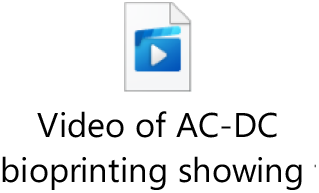
Video of AC-DC bioprinting showing the formation of a 3D implant. A 3D graft with eight layers of collagen fiber, 5 mm width, and 10 mm height is printed onto a temporary frame using the AC-DC process.

## Acknowledgments

The authors wish to thank P. Sharma, N. Kemper, and N. Sori for their contributions and efforts.

## Author contributions

KWC performed AC-DC bioprinting conceptualization and design, implant fabrication, cell viability/directionality/distribution analysis, mechanical property testing, data analysis related to *in vitro* analysis, and contributed to manuscript preparation. JT performed animal surgeries and care, functional testing, immunohistochemistry, SMASH analysis, data analysis related to *in vivo* analysis, and contributed to manuscript preparation. KC assisted with implant fabrication and *in vitro* analysis. YM and AB assisted with *in vitro* analysis. OS and MR assisted with developing SMASH protocols. AR assisted with SMASH analysis. AS contributed to manuscript preparation. GJC provided project guidance and contributed to manuscript preparation. MPF provided project guidance, AC-DC bioprinting conceptualization, and contributed to manuscript preparation.

## Data and materials availability

Materials described in this article may be available through material transfer agreements with the authors and their respective affiliations.

